# Variation in the modality of a yeast signaling pathway is mediated by a single regulator

**DOI:** 10.1101/131938

**Authors:** Julius Palme, Jue Wang, Michael Springer

## Abstract

Bimodal gene expression by genetically identical cells is a pervasive feature of signaling networks. In the galactose-utilization (GAL) pathway of *Saccharomyces cerevisiae*, induction can be unimodal or bimodal depending on natural genetic variation and pre-induction conditions. Here, we find that this variation of modality is regulated by an interplay between two features of the pathway response, the fraction of cells that are in the induced subpopulation and their expression level. Combined, the variations in these features are sufficient to explain the observed effects of natural variation and pre-induction conditions on the modality of induction in both mechanistic and phenomenological models. Both natural variation and pre-induction conditions act by modulating the expression and function of the galactose sensor *GAL3*. The ability to alter modality may allow organisms to adapt their level of “bet hedging” to the conditions they experience, and thus help optimize fitness in complex, fluctuating natural environments.

## Introduction

Non-genetic heterogeneity is a pervasive feature of gene expression and cellular signaling (1–3). Bimodal responses, where cells in an isogenic population adopt one of two distinct states, are particularly important for microbes coping with fluctuating environments (4, 5) and cells of multicellular organisms differentiating into discrete types (6, 7).

The galactose-utilization (GAL) pathway in *Saccharomyces cerevisiae* is a well-characterized bimodal response and a classic model of microbial decision-making (8, 9). Bimodality of GAL gene expression has been attributed to bistability arising from positive feedback through the Gal1p kinase and the Gal3p transducer (10, 11). Perturbing many of the components of the GAL pathway such as the Gal2p permease, the Gal4p activator, and the Gal80p repressor have been found to affect quantitative features of the GAL response (11–14) and in principle could affect the feedback in the system. However, only changes in Gal1p and Gal3p (10, 11) have been shown to affect modality, i.e. whether the response is bimodal or unimodal.

Our existing insight into modality in the GAL system comes almost entirely from measuring one pathway phenotype, the induced fraction, under one environmental perturbation, galactose titration (10, 13, 15– 17). The few studies that have deviated from this experimental approach have resulted in observations that raise new questions. For example, the GAL response was found to be unimodal or bimodal depending on the carbon source prior to encountering galactose (18); the molecular basis of this behavior is unknown. In our own previous work, we found that natural yeast isolates differ in the inducibility of GAL genes in mixtures of glucose and galactose (17, 19), and that some strains showed a bimodal response while others had a unimodal response (17). This natural variation provided an opportunity to dissect the genetic variants modulating bimodality in nature and gain insight into the quantitative behaviors organisms have evolved to respond to their environments.

In this work, we confirm and expand the observation that the pattern of GAL pathway induction can be either unimodal or bimodal depending on genetic background and pre-induction conditions. A computational model of GAL induction led us to a conceptual framework for variation in modality that identified the induced fraction and the threshold for glucose inhibition of expression as the two critical factors. Using this simple framework, we can explain the variations in modality we observed, and predict how new perturbations affect modality. Finally, we have determined that both natural variation and pre-induction conditions achieve changes in modality by tuning the expression and activity of a single signaling protein in the GAL pathway. These results reveal a simple evolutionary mechanism by which organisms can shape their responses to the environment, and suggest that modality is a highly adaptable feature of a signaling response.

## Results

### Genetic and environmental factors affect the modality of the GAL response

To study natural variation in the modality of the GAL response, we measured the expression of a *GAL1* promoter driving YFP (GAL1pr-YFP) in 34 geographically and ecologically diverse yeast strains (19–21) grown in different combinations of glucose and galactose (Figure 1A). When we titrated galactose concentration in the absence of glucose, all isolates showed a bimodal response with a mixture of uninduced and induced cells (Figure 1B, Figure S1). However, when we titrated glucose concentration in a constant concentration of galactose, the modality of natural isolates was variable: some still displayed bimodal population distributions (Fig 1C, ‘DBVPG1106’), while others displayed unimodal population distributions with intermediate *GAL1* expression (Figure 1C, ‘BC187’, Figure S2). We also found that growth history affected modality. For example, the laboratory strain S288C showed unimodal GAL induction when grown with raffinose as a carbon source prior to encountering mixtures of glucose and galactose, but gave a bimodal response when mannose was used as the initial carbon source instead (Figure 1D, (18)). These observations suggest that the modality of the GAL pathway is variable if glucose is present as well as galactose. However, it is unclear how adding a second input to a bimodal system can cause this qualitative change in pathway induction.

**Figure 1:**
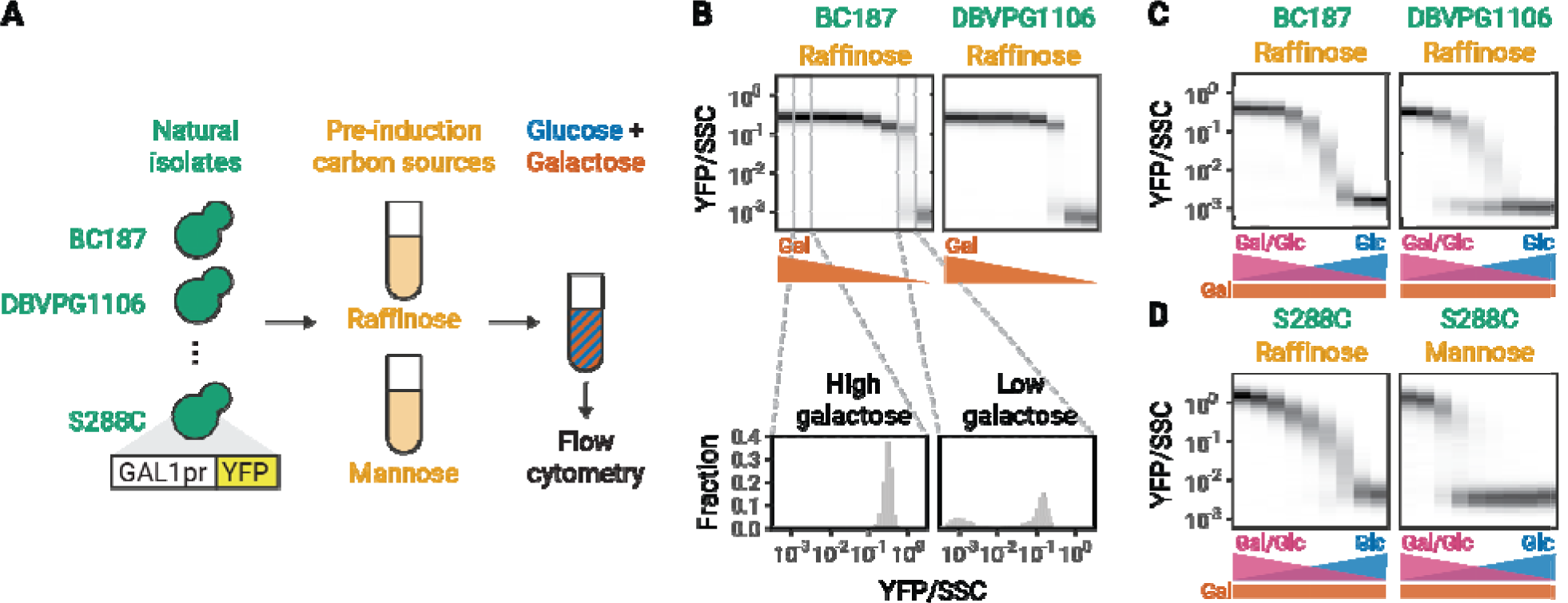
Genetic and environmental factors change GAL modality. (A) Experimental work-flow. Natural isolates of yeast tagged with a fluorescent reporter of *GAL1* expression were first grown in S medium with a pre-induction carbon source for 16 hours, then switched to S medium with mixtures of glucose and galactose. After 8 hours, *GAL1* expression was analyzed by flow cytometry. (B) GAL induction in different galactose concentrations. Bottom: Induction profiles of two natural isolates. Each plot represents 9 histograms with color intensities corresponding to the density of cells with a given value. Galactose concentration is titrated in two-fold steps from 1% to 0.0039%. Top: Blow out of two histograms of YFP level normalized by side scatter (SSC) at galactose concentrations 1% (left) and 0.0078% (right). (C) GAL induction of two natural isolates in mixtures of glucose and galactose. Glucose concentration was titrated in two-fold steps from 0.0039% to 1%. Galactose concentration was kept constant at 0.25%. (D) GAL induction of the laboratory strain S288C in mixtures of glucose and galactose after pre-induction growth in different carbon sources. Since the GAL pathway responds to the ratio of galactose and glucose (22), increasing the glucose concentration while keeping the galactose concentration constant simultaneously decreases GAL activation and increases glucose repression. All measurements are representative examples of at least two independent repeats.

### The interplay between GAL induction and expression level regulation determines modality

Bimodality in the GAL pathway is commonly attributed to bistability arising from positive feedback. To investigate the mechanism by which glucose regulates bistability (and thus the modality) of GAL induction, we adapted a previous ODE model that described the regulation of the GAL pathway by glucose and galactose (15). In this model, Gal80p binds the transcription factor Gal4p and keeps it in an inactive state. In the presence of galactose, Gal3p binds Gal80p, freeing Gal4p from the Gal80p-Gal4p complex. Bistability can arise in this system as a result of transcriptional feedback loops where free Gal4p leads to the production of Gal3p and Gal80p (10). Conversely, glucose inhibits GAL activation as it reduces the intracellular galactose concentration through competition for binding to hexose transporters (22) and activates Mig1p which in turn decreases the production of Gal3p and Gal4p (Figure 2A).

**Figure 2:**
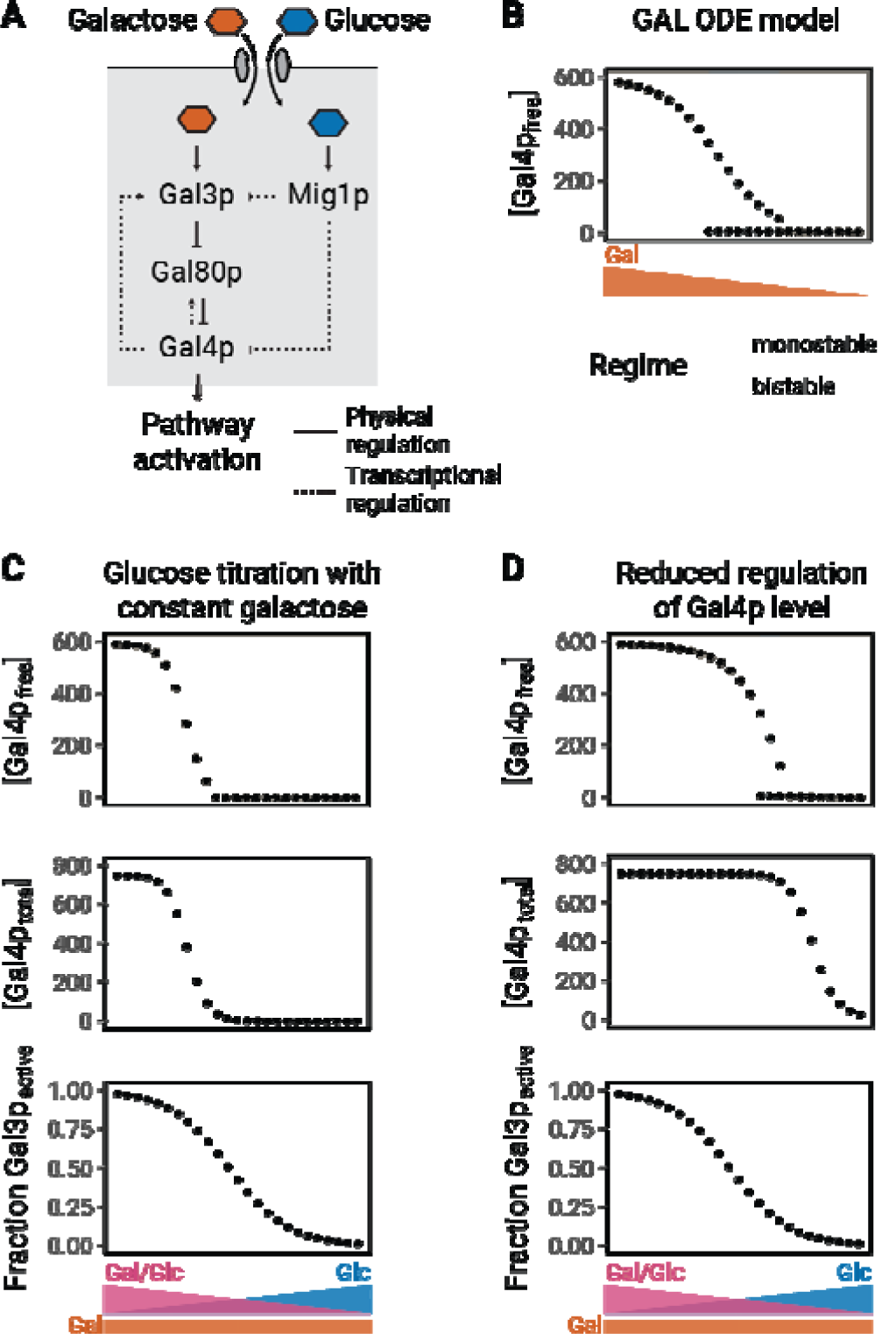
Glucose leads to a monostable response in an ODE model of the GAL pathway. (A) Schematic description of the ODE model. (B) Steady-state concentration of free Gal4p after simulations in different galactose concentrations in the absence of glucose. Regions are called bistable if the difference in steady-state free Gal4p concentration between two initial conditions is larger than 1. (C-D) Steady-state concentrations of free Gal4p, total Gal4p and the fraction of free Gal3 that is in its active state after simulations in different glucose concentrations with a constant background of galactose. Simulations were performed with (C) stronger or (D) weaker inhibition of Gal4p level by Mig1p (see Methods).

To analyze how glucose affects bistability in this model, we simulated the ODE system in different concentrations of glucose and galactose. Since Gal4p drives the expression of GAL genes, we used the steady-state concentration of free Gal4p as a read-out for pathway activity. To determine bistability in the system, we started all simulations from two extreme initial conditions, the steady-state concentrations of components in pure galactose or pure glucose. We then simulated switching to a mixture of sugars, and analyzed whether these initial conditions led to different steady-state levels of components in the new environment. As expected, the system displays bistability when galactose is titrated in the absence of glucose (Figure 2B). Next, we analyzed the effect of titrating glucose in the presence of a constant concentration of galactose, mirroring our experimental approach (Figure 1C, 1D). In the model, the addition of glucose converts the system from bistable to monostable (Figure 2C), indicating that the ODE system recapitulates the experimentally observed effect of glucose addition. To investigate the molecular basis for the switch, we then analyzed the effects of glucose in the simulations. Glucose has two effects: 1) it directly increases the activity of the transcriptional repressor Mig1p and 2) it indirectly decreases the intracellular concentration of galactose through competition for binding to transporters. Therefore, increasing glucose decreases both the total transcriptional output of *GAL3* and *GAL4* and the fraction of Gal3p that is in the active form (Figure 2C). The indirect effect of glucose on Gal3p activity is equivalent to the direct effect of galactose on Gal3p activity. As the system is bistable in pure galactose, this indirect effect cannot be the molecular cause of the change in bimodality; therefore, glucose must eliminate bistability through Mig1p.

Molecularly, bistability occurs when the system is near the activation threshold of Gal3p (10). Noise in the level of active Gal3p presumably leads to some cells to be above the activation threshold while others are below it. This difference is amplified by positive feedback, leading to a bimodal distribution. A second requirement is that Gal4p must be expressed at sufficient levels near the Gal3p activation threshold, so that Gal4p mediated positive feedback can occur. Figure 2C shows that when Gal4p abundance is low near this threshold, the system is monostable. To further test this hypothesis, we varied the Michaelis constants of parameters that affect glucose-dependent inhibition of Gal4p production (see Methods). These simulations confirmed that the system is bistable only when the concentration of Gal4p is sufficient to drive positive feedback upon activation and the system is near the Gal3p activation threshold (Figure 2D). In contrast, varying a parameter that controls the Mig1p inhibition of *GAL3* had no effect on whether the simulations were monostable or bistable (Figure S3).

While many unknown molecular parameters can affect the ratio of galactose and glucose required for Gal3p activation and the concentration of glucose required for Mig1p repression of Gal4p, systematic titrations of parameters that control these two processes show that modality depends on their relative thresholds (Figure S4). We hypothesized that a simpler phenomenological model focusing only on two free parameters should be able to recapitulate the variation in modality we experimentally observed. We therefore constructed a model in which the fraction of active Gal3p determines the fraction of cells that are in the induced state and the amount of free Gal4p determines the *GAL1* expression level of the induced subpopulation (Figure 3A). Mathematically, we described these two behaviors as Hill functions that decrease with increasing glucose concentration. To simulate induction profiles, we generated population distributions at different glucose concentrations by using the induced fraction to set the relative size of the two populations, the expression level to set the mean level of the induced subpopulation, and a normal distribution based on experimental noise to set the spread in expression. This simplified model focuses on how modality is affected by differences in the pathway’s responses to sugar ratio and glucose repression.

**Figure 3:**
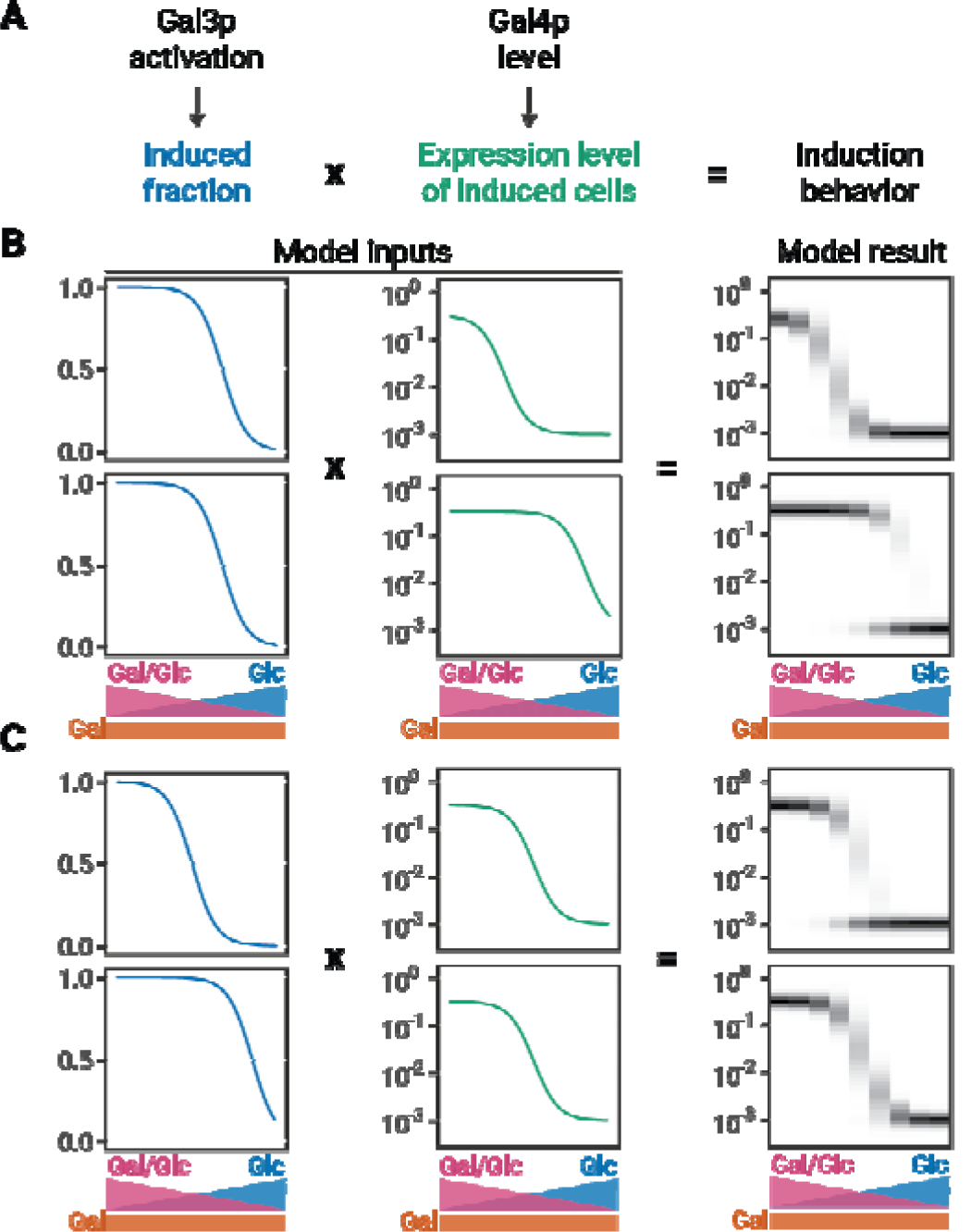
Phenomenological modeling of GAL induction. (A) The population distribution of GAL induction in every condition is derived based on the induced fraction (determined by the extent of Gal3p activation) and the expression level of induced cells (determined by the amount of free Gal4p). (B-C) Model results for varying expression level regulation with constant induced fraction regulation or varying induced fraction regulation with constant expression level regulation (C). The induced fraction and expression level of induced cells are described as functions that decrease with increasing glucose concentration. The induced fraction and induced level curves are used to sample uninduced and induced subpopulations from normal distributions. For the uninduced subpopulation, the relative expression level is 10^-3^. For the induced subpopulation, the expression level is given by the induced level curve. The relative size of the two populations is given by the induced fraction curve. Model results are represented by histograms at 9 different glucose and galactose combinations. The intensity of the color on the plot corresponds to the density of cells with a given induction value.

To confirm that the simplified model provides the same qualitative results as our molecular model, we varied the glucose threshold for the expression level of induced cells while keeping the glucose threshold for the induced fraction constant. In agreement with the simulations from our molecular model, the induction profile is unimodal when the induced fraction changes at concentrations of glucose that still inhibit expression level (Figure 3B, top). Conversely, the induction profile is bimodal when the induced fraction is in a regime of greatest change while the expression level is not inhibited by glucose (Figure 3B, bottom). Next, we varied the glucose threshold for the induced fraction while keeping the glucose threshold for the expression level constant. Again, these changes were sufficient to change modality (Figure 3C). Together, these results show that varying the relative position of induced fraction and expression level regulation in a phenomenological model is sufficient to recapitulate changes in modality.

We previously showed that induced fraction and expression level are regulated by the galactose/glucose ratio or the glucose concentration, respectively, and that natural genetic variation can change either of these parameters selectively (23). This raises the question of whether natural genetic variation can also alter GAL induction profiles along the axes that we identified in our mechanistic and phenomenological modeling. To analyze induced fraction and expression level regulation in our data, we computationally identified the induced and uninduced subpopulations by comparing GAL reporter distributions to the distribution of an uninduced sample (as in (16), Figure S5). We then calculated two summary metrics for each strain’s behavior: E_10_ (‘Expression level threshold’), the glucose concentration where the GAL1 expression level of the induced subpopulation reaches 10% of its level in pure galactose (Figure 4A), and F_90_ (‘induced Fraction threshold’), the glucose concentration where 90% of cells are in the induced subpopulation (Figure 4B).

**Figure 4:**
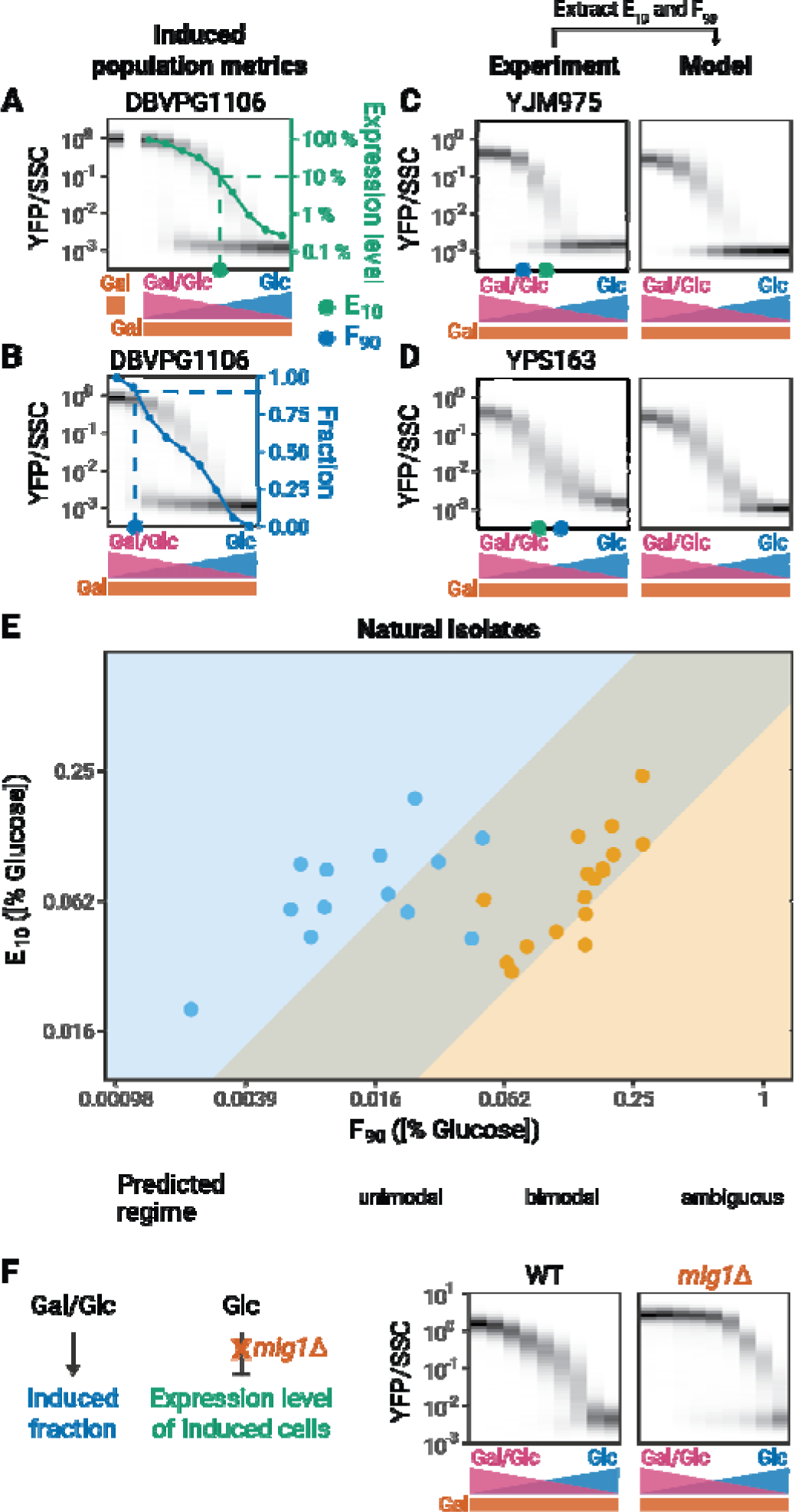
Experimental validation of model predictions. (A) Expression level metric (E_10_). GAL induction is measured in 2% galactose to determine the maximal expression level. The E_10_ is the glucose concentration at which the expression level of the induced subpopulation reaches 10% of the expression level in 2% galactose in the absence of glucose. (B) Induced fraction metric (F_90_). The induced fraction for each strain is determined by comparing the expression level of cells in the population with that of an uninduced population grown in 2% glucose, and calculating the fraction of cells with an expression level that is outside the range for the uninduced population (Figure S5). The F_90_ is the glucose concentration where the induced fraction reaches 90%. (C-D) Modality prediction based on induction metrics. In the phenomenological model, the position of the induced fraction and induced level functions are determined by the F_90_ and the E_10_, respectively. (E) F_90_ and E_10_ values of a panel of 30 natural isolates. Values correspond to the mean of 2-5 replicates. Modality was determined by comparing the fit of the data to a single or double Gaussian model (See Methods). Background colors correspond to predicted regimes for unimodal and bimodal strains. Simulations with diverse combinations of F_90_ and E_10_ values as well as different slopes for the induced fraction and induced level curves delineate regimes of unimodal and bimodal behaviors. The overlap represents an ambiguous regime where both unimodal and bimodal behaviors are possible. (F) Effect of *mig1*Δ on *GAL1* induction profiles.

Our modeling suggests that determining E_10_ and F_90_ should be sufficient to predict whether a strain is bimodal or unimodal. To test this hypothesis, we calculated the E_10_ and F_90_ values of natural isolates and used them as inputs to determine the position of the induced fraction and expression level functions in our phenomenological model. For both bimodal and unimodal strains, the experiments and models are in good agreement (Figure 4C, 4D, Figure S6, S7). A minor area of disagreement is that metrics from unimodal strains often predict a narrow range of bimodality in our model. We believe there are two factors that might explain these erroneous predictions. First, to measure the F_90_, there must be measurable gene induction. Therefore, the calculated F_90_ is a lower bound for the actual value of F_90_ in unimodal strains; in many cases the calculated F_90_ will be higher than the actual F_90_. If the calculated F_90_ for simulations of unimodal strains is increased by even a factor of 2, the predicted bimodal area disappears (Figure S8). Second, the slopes of the induced fraction curves could also be subject to variation and affect modality. Increasing the steepness of the induced fraction curve or induced level curves by increasing the Hill coefficients in our models can make simulations more unimodal (Figure S9). Indeed, some natural isolates appear to have steeper induction curves (Figure S10). These two explanations are not mutually exclusive, and we believe that these minor errors do not substantially detract from the value of our model.

We next scanned the parameters for our phenomenological model using a wide range of summary metrics and slopes to delimit a phase diagram of GAL induction modality. We then compared this phase diagram to the experimental data from natural isolates (Figure 4E). The modality of the natural isolates agrees well with their predicted modality in the phase diagram, supporting our hypothesis that the E_10_ and F_90_ measurements capture the important biological features that determine the modality of a strain.

Our ODE simulations gave us a molecular explanation for unimodality: Mig1p activity leads to unimodality by inhibiting *GAL4* expression in the regime where Gal3p activation is increasing. Therefore, an important prediction of the model is that removing Mig1p regulation should restore bimodality in unimodal strains. To test this prediction, we analyzed the induction profile of a *mig1*Δ strain. Deleting *MIG1* removes the glucose dependent regulation of *GAL4* expression level and thus the E_10_ of the deletion strain is increased compared to the wild-type strain (Figure 4F). As predicted, it also converts the strain from unimodal to bimodal. In addition, the observed F_90_ value for the bimodal *mig1*Δ strain is higher than that for the unimodal wild-type. This supports our hypothesis that Mig1p-dependent repression conceals the actual F_90_ value of unimodal strains and that the observed F_90_ value of unimodal strains is only a lower bound for the actual F_90_ value.

### History-dependence of modality can be explained by changes in the induced fraction

It has previously been reported that pre-induction growth conditions can affect the modality of GAL induction (18), which offers another opportunity to test the predictions from our modeling framework. To see how metabolic history affects F_90_ and E_10_, we grew 13 natural isolate strains in mannose, raffinose, acetate, or glycerol prior to transferring them into mixtures of glucose and galactose (Figure S11). There are three broad categories of strain responses to pre-induction conditions. The first category contains strains that are unimodal when pre-grown in raffinose that become bimodal when pre-grown in mannose (Figure 5A, 5B). The second is strains that are bimodal when pre-grown in raffinose but become unimodal when pre-grown in glycerol or acetate (Figure 5C, 5D). The last is strains that are always bimodal or always unimodal in all pre-induction carbon sources tested here (Figure 5E, 5F). Analyzing the effect of pre-induction conditions on our summary metrics, we found that the average fold change between the lowest and highest F_90_ values of each strain is 8.61 while the average fold change for the E_10_ is 1.78 (Figure S12). Our heuristic model predicts that the magnitude of the changes in F_90_ values are sufficient to explain the observed changes of modality. Based on this analysis, the modality of all our experimental results agrees with their position in our phase diagram of GAL induction (Figure 5B, 5D, 5F).

**Figure 5:**
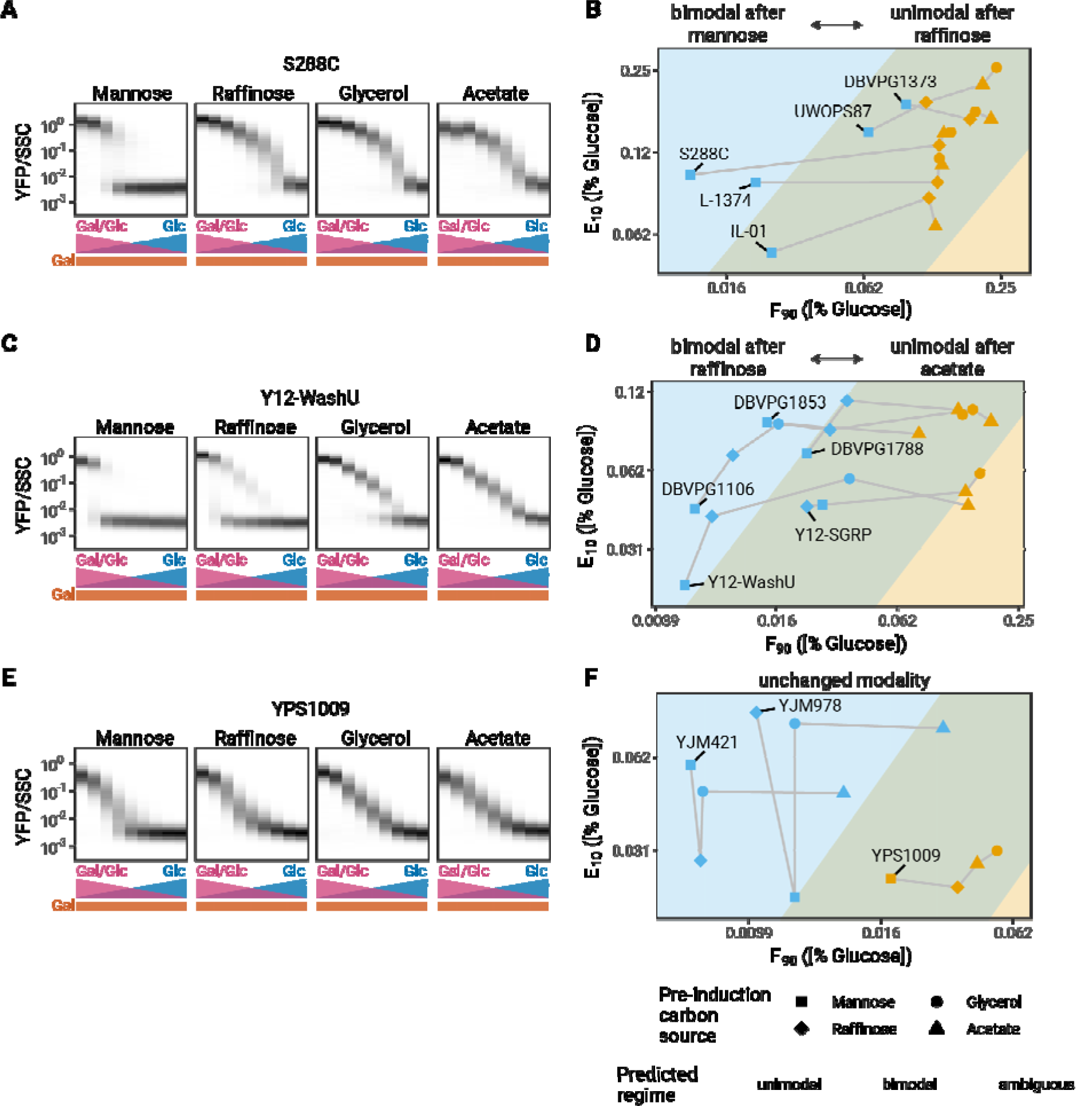
Metabolic history changes modality. (A) History-dependence of induction profiles. Induction profiles of S288C in mixtures of glucose and galactose after pre-induction growth in different carbon sources for 16 hours. (B) F_90_ and E_10_ values for isolates that are unimodal after growth in raffinose and bimodal after growth in mannose. (C) Induction profiles of Y12-WashU after growth in different carbon sources for 16 hours. (D) F_90_ and E_10_ values for isolates that are bimodal after growth in raffinose and unimodal after growth in either acetate or glycerol. (E) Induction profiles of YPS1009 after growth in different carbon sources for 16 hours. (F) F_90_ and E_10_ values for isolates that are always unimodal or bimodal. All measurements are representative examples of at least two independent repeats.

### Regulation of *GAL3* expression explains the history-dependence of the GAL response

To determine how pre-induction carbon source modulates F_90_, we measured the expression of GAL genes in pre-induction conditions using transcriptional reporters (Figure 6A). We found that GAL genes are down-regulated in carbon sources that lead to bimodal induction (mannose) and up-regulated in carbon sources that lead to unimodal induction (acetate, glycerol). Amongst all GAL genes, the expression levels of *GAL3* and *GAL4* show the strongest fold change between the carbon sources tested (Figure 6A).

**Figure 6:**
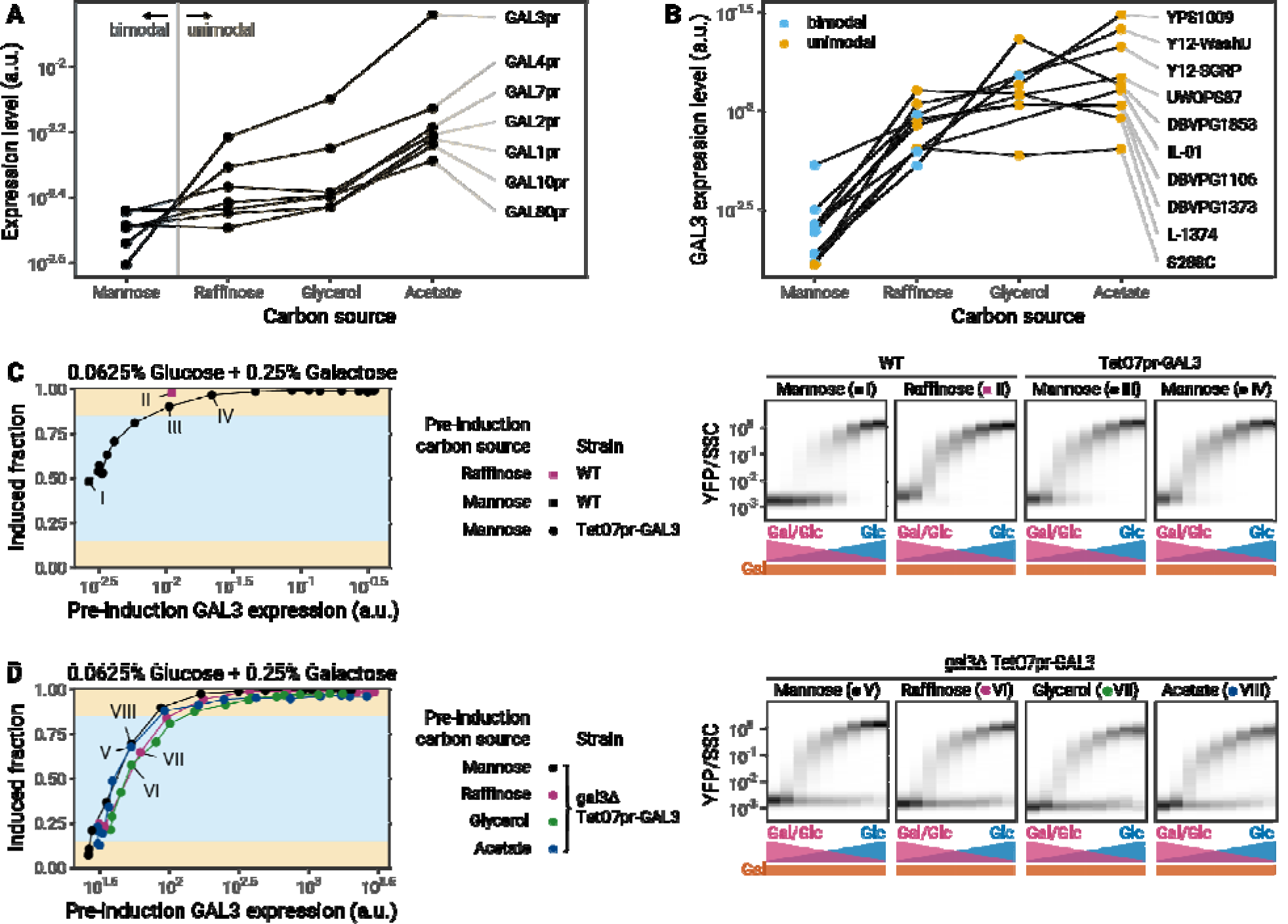
Pre-induction *GAL3* levels determine the modality of induction profiles. (A) Expression of GAL genes in different pre-induction carbon sources in S288C as determined by fluorescence of GAL promoter-YFP protein transcriptional reporter strains. (B) Effect of pre-induction growth in different carbon sources on the expression of a *GAL3*^S288C^ reporter in S288C and 9 natural isolates. Colors correspond to the modality of the induction profile of these strains in the given pre-induction carbon source. (C) Left: Effect of *GAL3* overexpression during pre-induction growth in mannose. Connected points correspond to a doxycycline titration series for a S288C TetO7pr-*GAL3* strain. Right: Complete induction profiles of S288C in mannose (I) or raffinose (II) and S288C TetO7pr-*GAL3* in mannose at two concentrations of doxycycline that lead to *GAL3* expression levels that bracket the expression level of GAL3 from a raffinose pre-induction culture (III, IV). (D) Left: Effect of synthetic *GAL3* expression in a Δ*gal3* during pre-induction growth in different carbon sources. Connected points correspond to a doxycycline titration series in different carbon sources for a S288C Δ*gal3* TetO7pr-GAL3-mScarlet strain. Right: Complete induction profiles of S288C Δ*gal3* TetO7pr-GAL3-mScarlet after pre-induction growth in different carbon sources with constant *GAL3* expression.

We hypothesized that *GAL3* was more likely than *GAL4* to be the dominant factor due to its high dynamic range of expression (Figure 6A) and prior evidence for *GAL3* concentration affecting the GAL decision (13). We therefore analyzed the regulation of *GAL3* by a range of pre-induction carbon sources in 9 natural isolates using a transcriptional reporter for the *GAL3*^S288C^ promoter. We found that in each strain the *GAL3* expression level in a pre-induction conditions generally correlates with the modality observed later (Figure 6B).

To test if *GAL3* expression prior to induction is the key determinant of modality, we used a tetracycline-inducible promoter to control the expression of *GAL3* directly. We predicted that forcing a change in *GAL3* expression while keeping the pre-induction carbon the same should change modality. Conversely, changing the pre-induction carbon without changing *GAL3* expression should not change modality.

For the laboratory strain S288C, pre-induction growth in mannose leads to low *GAL3* expression and a bimodal induction profile, while pre-induction growth in raffinose leads to higher *GAL3* expression and a unimodal induction profile (Figure 5A, 6A). When we overexpressed *GAL3* during pre-induction growth in mannose using tetracyclin induction, we saw an increase in the induced fraction and a loss of bimodality (Figure 6C). Thus, the *GAL3* concentration pre-induction is sufficient to set the modality in this strain background. Artificially setting the *GAL3* level of mannose pre-induction cultures to that of a raffinose pre-induction culture converted the induction profiles to one similar to a raffinose pre-induction culture (Figure 6C, II and IV).

In addition to showing unimodal induction after raffinose pre-induction, this strain also shows unimodal behavior after pre-induction in acetate or glycerol (Fig. 5A). We deleted the endogenous *GAL3* gene from our tetracycline induction strain, so that we could force pre-induction *GAL3* expression to remain low in these sugars. As predicted, when pre-induction *GAL3* expression in these sugars was low we saw bimodal induction almost identical to that seen with mannose pre-induction (Fig. 6D). Pre-induction *GAL3* concentrations also set the induced fraction with almost no dependence on the pre-induction carbon source (Figure 6D). We conclude that regulation of *GAL3* expression in pre-induction conditions is the major driver of history-dependence in the modality of GAL induction.

### Natural variation in *GAL3* alleles changes the modality of induction by tuning the induced fraction

The central role of *GAL3* expression in setting the modality of induction suggests that natural variation in *GAL3* alleles could be responsible for the observed differences in modality between isolates (Figure 1C). Previously, we showed that polymorphisms in the *GAL3* gene explain most of the natural variation in the decision to induce the GAL pathway (17), i.e. the F_90_ (17), suggesting that allele swaps of the *GAL3* ORF should alter the F_90_ of the strain, which in some cases would be enough to switch the modality of induction. To test this prediction, we determined the modality of a set of 30 allele swap strains comprised of 10 *GAL3* alleles in 3 genetic backgrounds. The experimentally determined modality of all allele swap strains agrees with the expected modality in each case (Figure 7A, Figure S13). For example, replacing the *GAL3* allele of two unimodal strains, BC187 and S288C, with the *GAL3* allele of any of our bimodal strains is sufficient to change the induction profiles from unimodal to bimodal (Figure 7B, 7C). In agreement with our model, the F_90_ in these strains decreases sufficiently to convert strains from unimodal to bimodal. Replacing the *GAL3* gene of the bimodal strain YJM978 with the alleles of unimodal strains does not change the modality (Figure 7D). This is also in agreement with our model which predicts that the magnitude of the increase in F_90_ caused by allele swaps in the YJM978 strain background are insufficient to change modality (Figure 7D).

**Figure 7.**
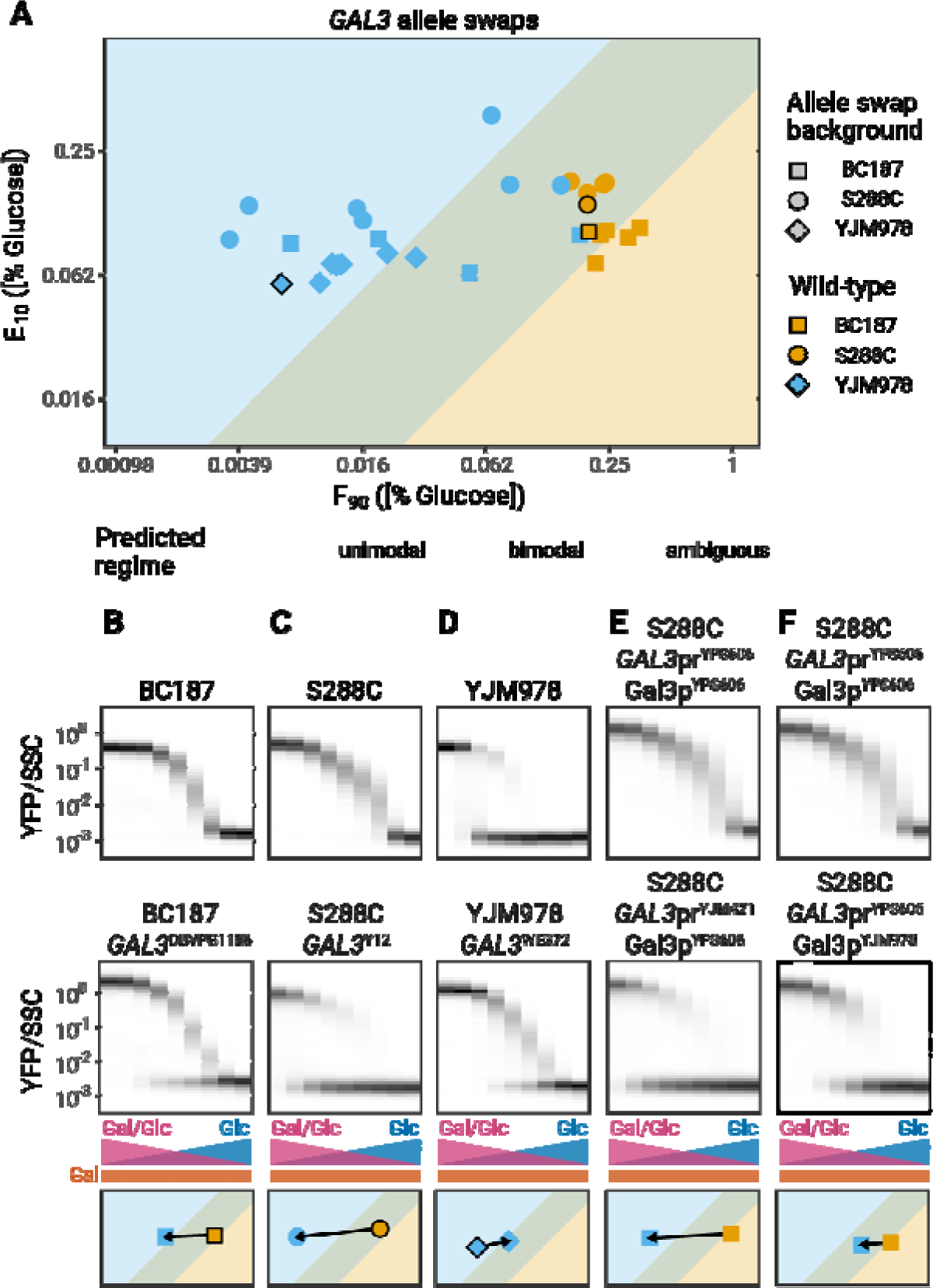
Allele swaps of *GAL3* change modality. (A) F_90_ and E_10_ values for a panel of allele swaps (10 *GAL3* alleles in 3 different genetic backgrounds). Strains that never reach an induced fraction of 90% are not shown here. Black outlines denote the wild-type strains. All measurements are representative examples of at least two independent repeats. (B-F) Effect of *GAL3* allele swaps on modality. (Top) Induction profiles of (B-D) wild-type isolates or (E-F) S288C *GAL3*^YPS606^ (Middle) Induction profiles of (A-C) *GAL3* allele swaps, (D) *GAL3* promoter swaps or (F) *GAL3* CDS swaps. (Bottom) Effect of the perturbation on F_90_ and E_10_. Arrows start at the values of the wild-type or S288C *GAL3*^YPS606^ strains and end at the values of the perturbed strain.

To further explore the variations in modality amongst the natural isolates, we analyzed the contribution of promoter and coding sequence variation. We found that SNPs in either promoter or coding regions (CDS) are sufficient to change modality (Figure 7E-F). For example, S288C with *GAL3*^YPS606^ is unimodal, but replacing the YPS606 promoter in this strain with the YJM421 promoter leads to bimodal induction (Figure 7E). Similarly, BC187 with *GAL3*^NC-02^ is unimodal, but replacing the NC-02 coding sequence with the YJM978 coding sequence leads to bimodal induction (Figure 7F). The mechanisms by which promoter and CDS changes are able to change the modality will be the subject of future work.

## Discussion

In mixtures of glucose and galactose, the response of the GAL pathway in a population of yeast cells can be either unimodal or bimodal depending on their evolutionary history and their current environmental conditions. Here, we show that the modality of GAL induction in different strains depends on the relative position of the sugar concentration thresholds at which induced fraction and expression level are regulated. Glucose inhibits the expression of GAL genes, preventing the positive feedback that is crucial for bistability. In general, any input to a pathway that can create or eliminate feedback has the potential to modulate bistability. Since many signaling responses are controlled by multiple inputs, our findings imply that other unimodal responses could be bimodal in different conditions and vice versa.

Bimodality in the GAL response is considered a bet-hedging strategy where a fraction of the population prepares for glucose depletion by inducing the GAL pathway while other cells maximize their current growth and do not induce the pathway (15, 19). This heterogeneity helps populations deal with uncertain, fluctuating environments. Bet-hedging is advantageous in the GAL system when the switching rate between glucose and galactose environments is high (24); indeed, cells evolve bimodality in MAL gene expression when they are continuously switched between glucose and maltose (25). Because cells can sometimes be in environments with a high switching rate and other times in environments with a low switching rate, a strategy that allows the extent of bet-hedging to be tuned could be the most optimal. In this work, we show strains can tune the amount of bimodality both physiologically, based on their metabolic history, and genetically, presumably based on the environmental statistics that different natural isolates have faced in their evolutionary history. Further work will be needed to determine the evolutionary consequences of tunable bimodal responses such as the ones we characterize here.

Previous work on cell-to-cell heterogeneity has typically emphasized the complex genetic architecture of the pathways involved (26, 27). In contrast, the physiological and genetic variation in modality in the GAL pathway can be explained by changes in the behavior of a single protein. Modulating levels of Gal3p, the intracellular galactose sensor, is sufficient for changing modality. Swapping the *GAL3* alleles of natural isolates can turn a unimodal strain into a bimodal strain. We show that the environment tunes the expression level of *GAL3*, and this tuning is sufficient to change the modality of GAL induction (Figure 7 and Figure S13). Circuit designs such as these, where a single gene controls modality, may have been selected in evolution, since they allow cells to easily adapt their behavior on both physiological and evolutionary timescales.

The control of GAL pathway modality by mannose, raffinose, glycerol, and acetate suggests an addition layer of metabolic regulation that has been largely missed in previous analyses of this pathway. These findings show that factors other than canonical glucose catabolite repression can be important in determining the inducibility of GAL genes, consistent with our findings that many mutants outside the GAL pathway can have a significant effect on GAL response (28). The fact that pre-induction carbon sources mostly affect F_90_, just as *GAL3* allele swaps do, suggests that the *GAL3* positive feedback loop may be a nexus of regulation of GAL genes by multiple signals in the cell. In future studies, understanding the metabolic regulation of this well-studied system could give insight into the connections between metabolism and metabolic signaling in a variety of systems.

## Methods

### Strains and media

Strains were obtained as described in (17, 19). An initial set of 36 strains were assayed in a glucose gradient (1% to 0.0039%) with a constant background of 0.25% galactose. Strains DBVPG6765, CLIB324, L-1528, M22, W303, YIIC17-E5 were excluded from downstream analysis due to poor growth in our media conditions. Strain 378604X was also excluded due to a high basal expression phenotype that was an outlier in our collection. The genetic basis of this behavior is likely an interesting topic for follow-up studies. All experiments were performed in synthetic minimal medium (“S”), which contains 1.7g/L Yeast Nitrogen Base (YNB) (BD, Franklin Lakes, NJ, USA) and 5g/L ammonium sulfate (EMD). In addition; D-glucose (EMD, Darmstadt, Germany), D-galactose (MilliporeSigma, St. Louis, MO, USA), mannose (MilliporeSigma), glycerol (EMD), acetate (MilliporeSigma), and/or raffinose (MilliporeSigma) were added as a carbon source. Cultures were grown in a humidified incubator (Infors Multitron, Bottmingen, CH) at 30°C with rotary shaking 999 rpm (500 uL cultures in 1 mL 96-well plates).

### Flow cytometry

Cells were struck onto YPD agar from −80C glycerol stocks, grown to colonies, then inoculated from colony into YPD liquid and cultured for 16-24 hours. Then, cultures were inoculated in a dilution series (1:200 to 1:6400) in S + 2% pre-induction carbon source medium. The pre-induction cultures were incubated for 16-20 hours, and then their optical density (OD600) was measured on a plate reader (PerkinElmer Envision). The outgrowth culture with OD600 closest to 0.1 was selected for each strain, and then washed twice in S (with no carbon sources). To determine expression levels in pre-induction conditions, washed cells were then diluted in Tris-EDTA pH 8.0 (TE) in a shallow microtiter plate (CELLTREAT, Pepperell, MA, USA). For sugar gradient experiments, washed cells were diluted 1:200 into the appropriate sugar in 96-well plates (500 uL cultures in each well) and incubated for 8 hours. Then, cells were harvested and fixed by washing twice in TE and resuspended in TE before transferring to microtiter plate for measurement. Flow cytometry was performed using a Stratedigm S1000EX with A700 automated plate handling system.

### *GAL3* titration in pre-induction conditions

To titrate *GAL3* levels in the presence of the native *GAL3* gene, the *AGA1* gene was replaced with a MYO2pr-rtTA-TetO7pr-GAL3 construct in a hoΔ:GAL1pr-YFP strain. Cells were grown for 16 hours in S + 2% mannose as described above, but the medium was supplemented with doxycycline (MilliporeSigma) concentrations ranging from 38.9 μg/ml to 0.0176 μg/ml in 1.5x dilutions steps. To measure the total *GAL3* expression level after pre-induction growth, the *AGA1* gene was replaced with a MYO2pr-rtTA-TetO7pr-YFP construct in a hoΔ:GAL3pr-YFP reporter strain. After pre-induction growth in the same dilution doxycycline concentrations, cells were harvested and YFP levels were determined using flow cytometry as described above.

To titrate *GAL3* levels in the absence of the native *GAL3* gene, the *AGA1* gene was replaced with a MYO2pr-rtTA-TetO7pr-GAL3-mScarlet construct in a gal3Δ hoΔ:GAL1pr-YFP strain. Cells were grown for 16 hours in S + 2% pre-induction carbon source as described above, but the medium was supplemented with doxycycline concentrations ranging from 38.9 μg/ml to 0.0176 μg/ml in 1.5x dilutions steps. To measure the total *GAL3* expression level after pre-induction growth, cells were washed and mScarlet levels were determined by fluorescence microscopy using a Hamamatsu Orca-R2 camera (Hamamatsu, Japan) on a Ti Eclipse inverted Nikon microscope (Tokyo, Japan). Microscopy images were analyzed using U-net (29) and custom Python scripts.

### Data analysis

Data analysis was performed using custom MATLAB scripts, including Flow-Cytometry-Toolkit (https://github.com/springerlab/Flow-Cytometry-Toolkit).

To determine the modality of GAL induction experiments, a Gaussian function was fitted to the population distribution for each of the 9 sugar combinations. If the degree-of-freedom adjusted R^2^ of the fit was less than 0.99, two Gaussian functions were fitted to the data. Distributions were then determined to be bimodal if the distance between the means of the Gaussians was more than twice of the highest standard deviation of the Gaussian (as in (10)) and the fraction of the smaller Gaussian was higher than 0.15. GAL induction experiments or simulations that had a bimodal distribution in at least one combination of glucose and galactose in all replicates were called bimodal.

### ODE model

The ODE system from (15) was implemented using the G1 term to describe Gal3p and the R term to describe Mig1p with the parameters described in Supplementary Table 1. To determine the intracellular galactose concentration (α_gal_), the extracellular galactose concentration was divided by the extracellular glucose concentration and multiplied with a conversion factor μ. Steady-state species levels were obtained by simulating the system using the ode function of the R package deSolve in different levels of galactose (logarithmically spaced between 10^0^ to 10^4^) and glucose (logarithmically spaced between 10^-2^ to 10^1.5^ with a constant galactose level of 500). To decrease the strength of Mig1p inhibition on Gal1p and Gal4p levels, the K_R1_ or K_R4_ parameters were multiplied by 50.

### Phenomenological model

Induction profiles were simulated from E_10_ and F_90_ metrics using functions that describe the induced fraction and the mean expression level of the induced subpopulations as a function of the glucose concentration. For the induced level, the following function was used:

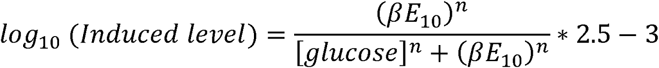

where 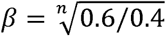 converts the glucose concentration where 10% of the maximal expression level is reached (i.e. the E_10_) to the glucose concentration where 50% of the maximal expression level is reached. The function was scaled from −3 to −0.5 to match the range of the experimental data. To obtain realistic versions for the n constant, this function was fitted to the induced level curves of natural isolates, the mean fitted n value was extracted for every natural isolate, and the mean of these values was used for simulations (induced level curve: 1.15, induced fraction curve: 1.69, see Figure S11).

For the induced fraction, the following function was used:

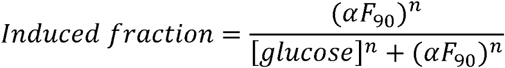

where the 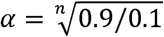 converts the glucose concentration where 90% of the cells are induced (i.e. the F_90_) to the glucose concentration where 50% of the cells are induced. This function was fitted to the induced level curves of bimodal natural isolates, the mean fitted n value extracted for every natural isolate, and the mean of these values was used for simulations (Figure S11).

For 9 concentrations of glucose and galactose, induced level and induced fraction values were extracted from these curves to generate simulated populations in these conditions. For a total population size of 20000 cells, uninduced and induced subpopulations were generated according to the induced fraction value. The expression level values of cells were drawn from normal distributions with the mean expression level of the uninduced subpopulation at a constant level of 10^-3^ and the level of the induced subpopulation at a value that was determined by the induced level curve. The standard deviations of the distributions were determined by fitting a quadratic equation to experimental standard deviations at different expression levels:

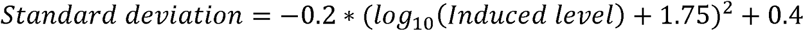

To delineate possible unimodal and bimodal regimes, GAL induction was simulated using all possible combinations of 10 different values for E_10_ and F_90_ (1, 0.5, 0.25, 0.125, 0.0625, 0.0313, 0.156, 0.0078, 0.0039, 0.0020, 0.0010). The Hill constants n for the induced fraction and the induced level functions in these simulations were varied between the lowest and highest experimentally fitted n values (induced level curve: 0.84 and 1.50, induced fraction curve: 0.75 and 2.95, see Figure S11). The E_10_ metric, the F_90_ metrics, and the modality of the induction profile were determined from these simulations as described above. In the F_90_-E_10_ space, unimodal and bimodal regimes were delineated by the bounding line with a slope of 1 that would capture all the unimodal or bimodal simulations respectively on one side of the line.

### Data availability

Raw flow cytometry and microscopy data have been deposited in Dryad (DOI: 10.5061/dryad.69p8cz8z8).

## Supplementary Information

**Supplementary Table 1.** Parameter values used in the ODE model.

| Parameter | Value |
| --- | --- |
| $\alpha_{0G1}$ | 4.25 |
| $\alpha_{0G80}$ | 7.1 |
| $\alpha_{G1}$ | 66 |
| $\alpha_{G4}$ | 3.75 |
| $\alpha_{G80}$ | 13.75 |
| $\alpha_R$ | 0.75 |
| $\gamma_{C81}$ | 0.005 |
| $\gamma_{C84}$ | 0.005 |
| $\gamma_{G1}$ | 0.005 |
| $\gamma_{G1s}$ | 0.005 |
| $\gamma_{G4}$ | 0.005 |
| $\gamma_{G80}$ | 0.005 |
| $\gamma_R$ | 0.005 |
| $\gamma_{Rs}$ | 0.005 |
| $k_{f81}$ | 82.2 |
| $k_{f84}$ | 95.2 |
| $k_{fg}$ | 562.4 |
| $k_{fR}$ | 112.4 |
| $K_{G1}$ | 25 |
| $K_{G80}$ | 5 |
| $K_{R1}$ | 0.5 |
| $K_{R4}$ | 0.5 |
| $k_{r81}$ | 700.1 |
| $k_{r84}$ | 1237 |
| $k_{rg}$ | 3391 |
| $k_{rR}$ | 3564 |
| $n_{80}$ | 3 |
| $n_{R1}$ | 1 |
| $n_{R4}$ | 3 |
| $\mu$ | 0.005 |

**Figure S1.**
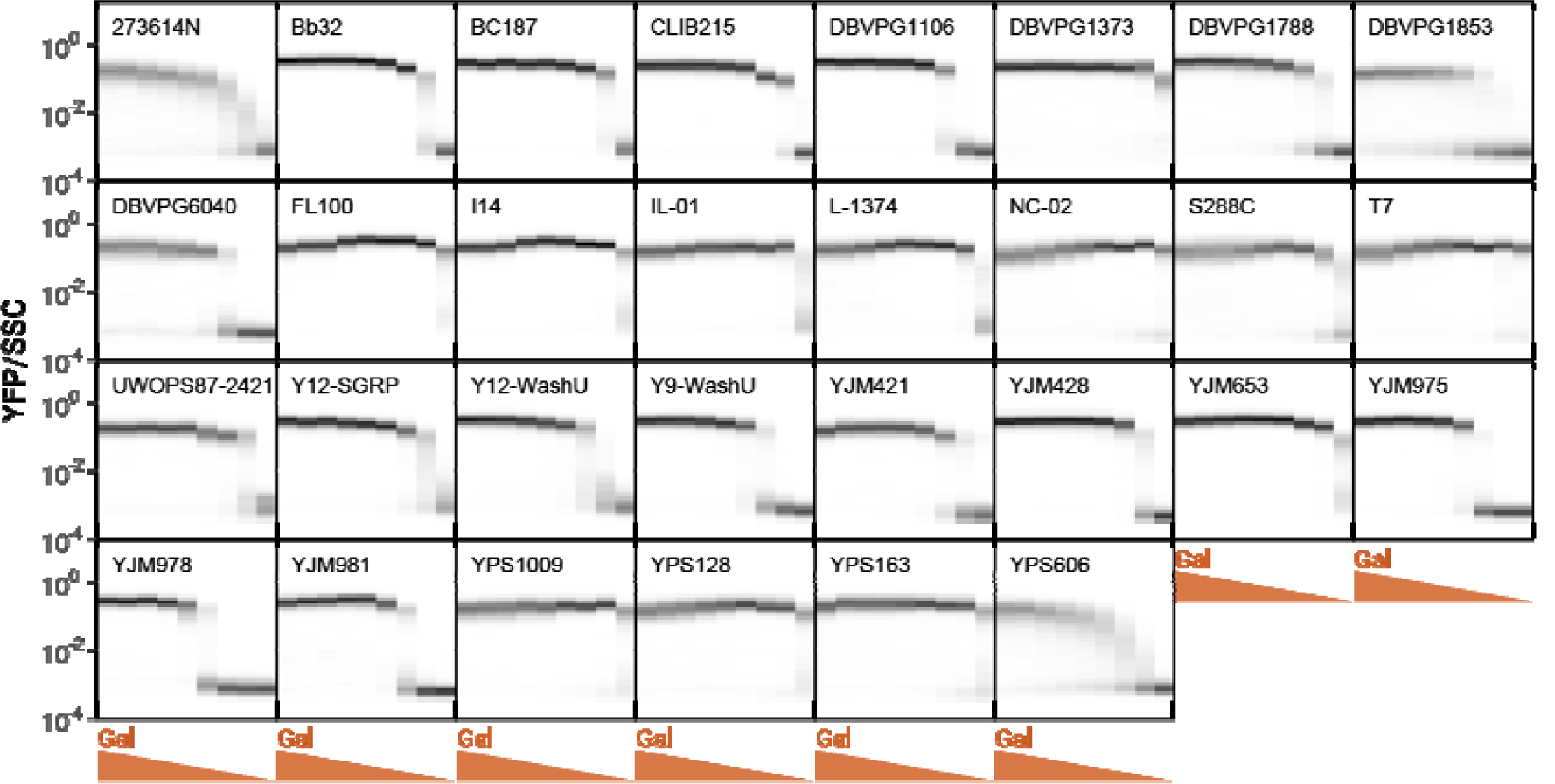
GAL induction of natural isolates in different galactose concentrations. Each plot represents 9 histograms with color intensities corresponding to the density of cells with a given fluorescence value (normalized by side-scatter (SSC)). Galactose concentration is titrated in two-fold steps from 1% to 0.0039%.

**Figure S2.**
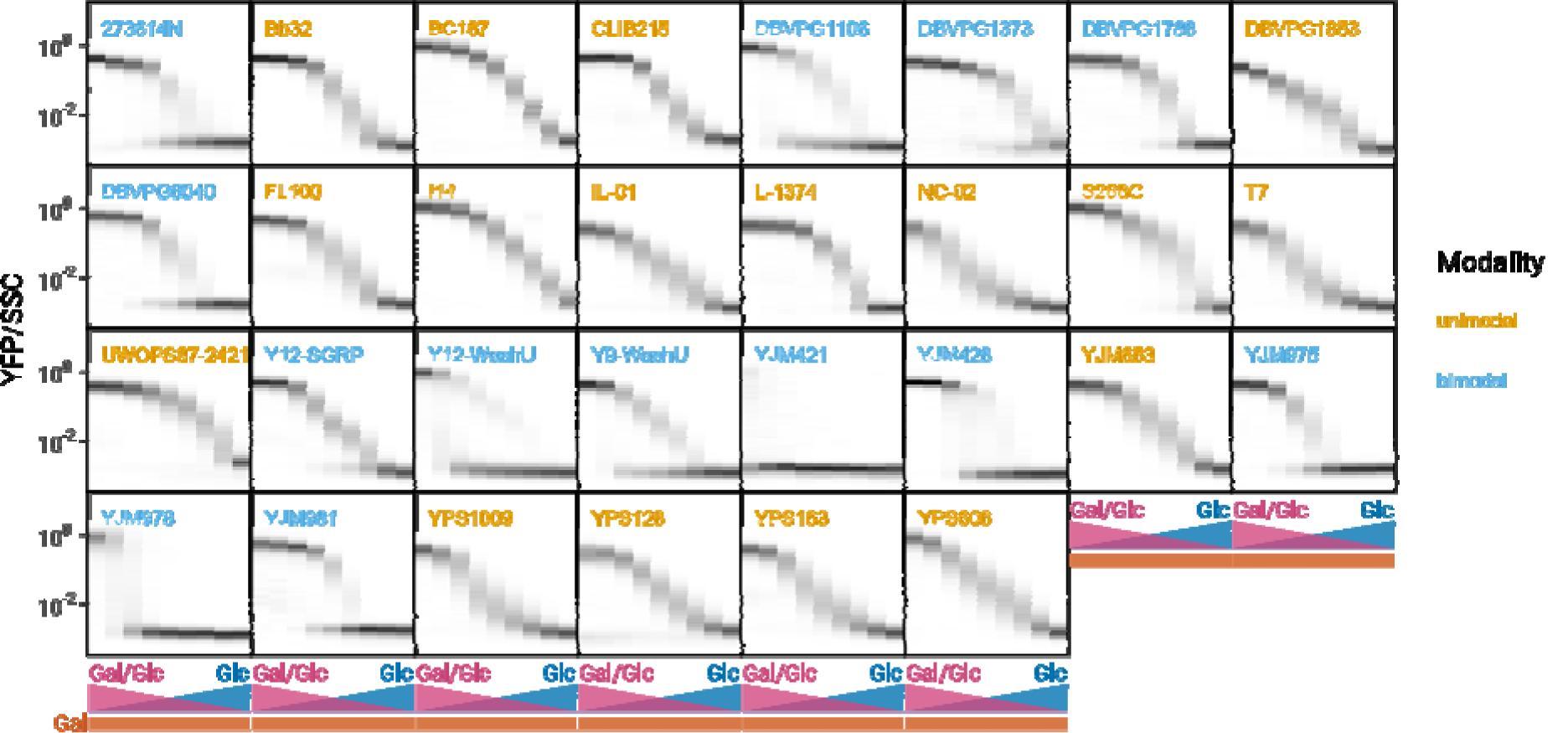
GAL induction of natural isolates in different glucose and galactose concentrations. Each plot represents 9 histograms with color intensities corresponding to the density of cells with a given fluorescence value (normalized by side-scatter (SSC)). Glucose concentration is titrated in two-fold steps from 1% to 0.0039%, galactose concentration is constant at 0.25%. Orange isolate names indicate unimodal induction, blue isolate names indicate bimodal induction.

**Figure S3.**
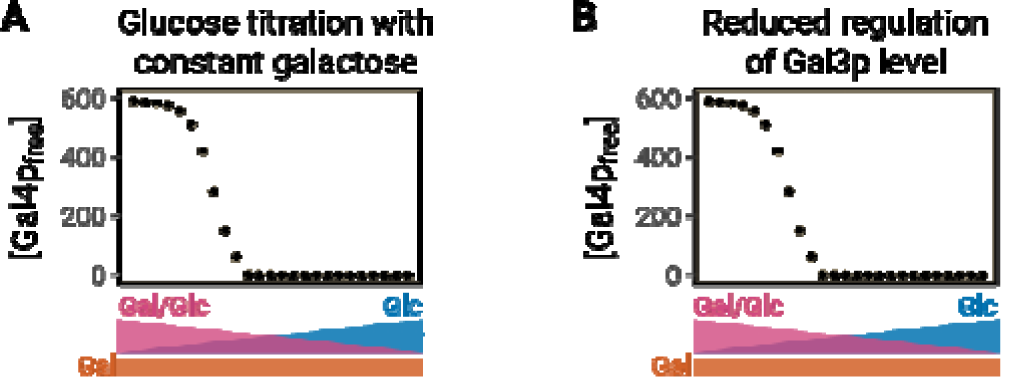
(A-B) Steady-state concentrations of free Gal4p after simulations in different glucose concentrations with a constant background of galactose. Simulations were performed with unchanged (A) or reduced (B) inhibition of Gal3p level by Mig1p.

**Figure S4.**
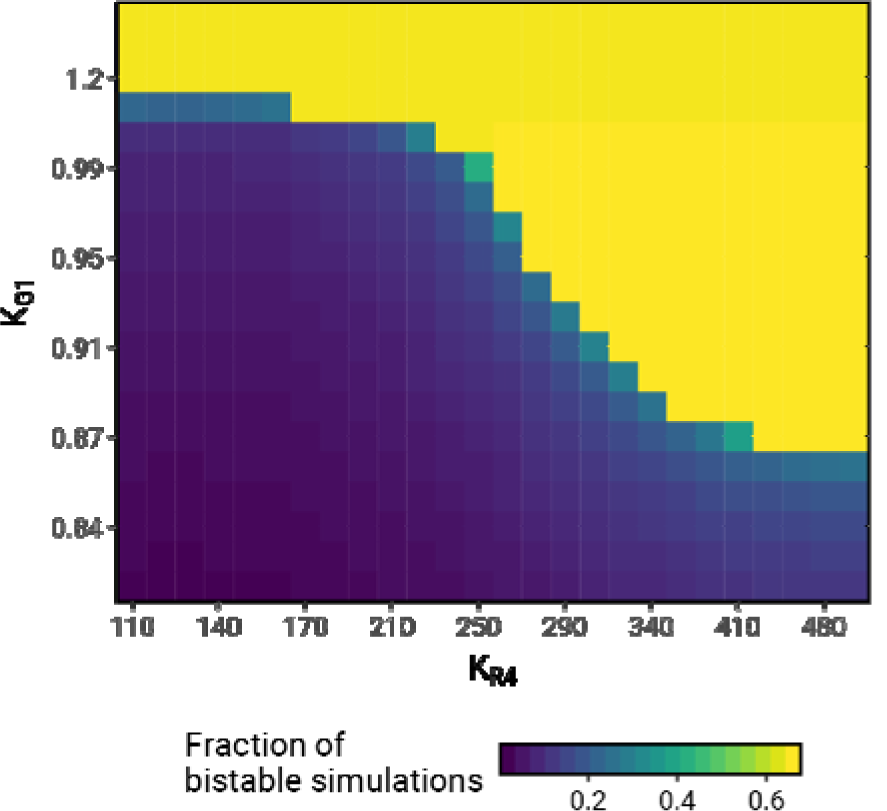
Phase space of glucose repression (K_R4_) and GAL induction (K_G1_) values. Parameters were multiplied with the indicated values before determining steady-state values in the same range of glucose and galactose values as in Figure 2, Figure S3. Bistability was assessed as described in Figure 2. The color indicates the fraction of glucose concentrations that showed bistability.

**Figure S5.**
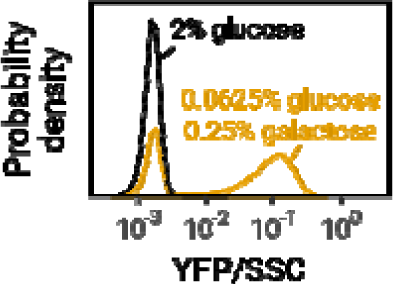
Identification of the induced subpopulation. A reference distribution from 2% glucose (black histogram) is subtracted from the population distribution (orange histogram) to yield the induced subpopulation (orange shading).

**Figure S6.**
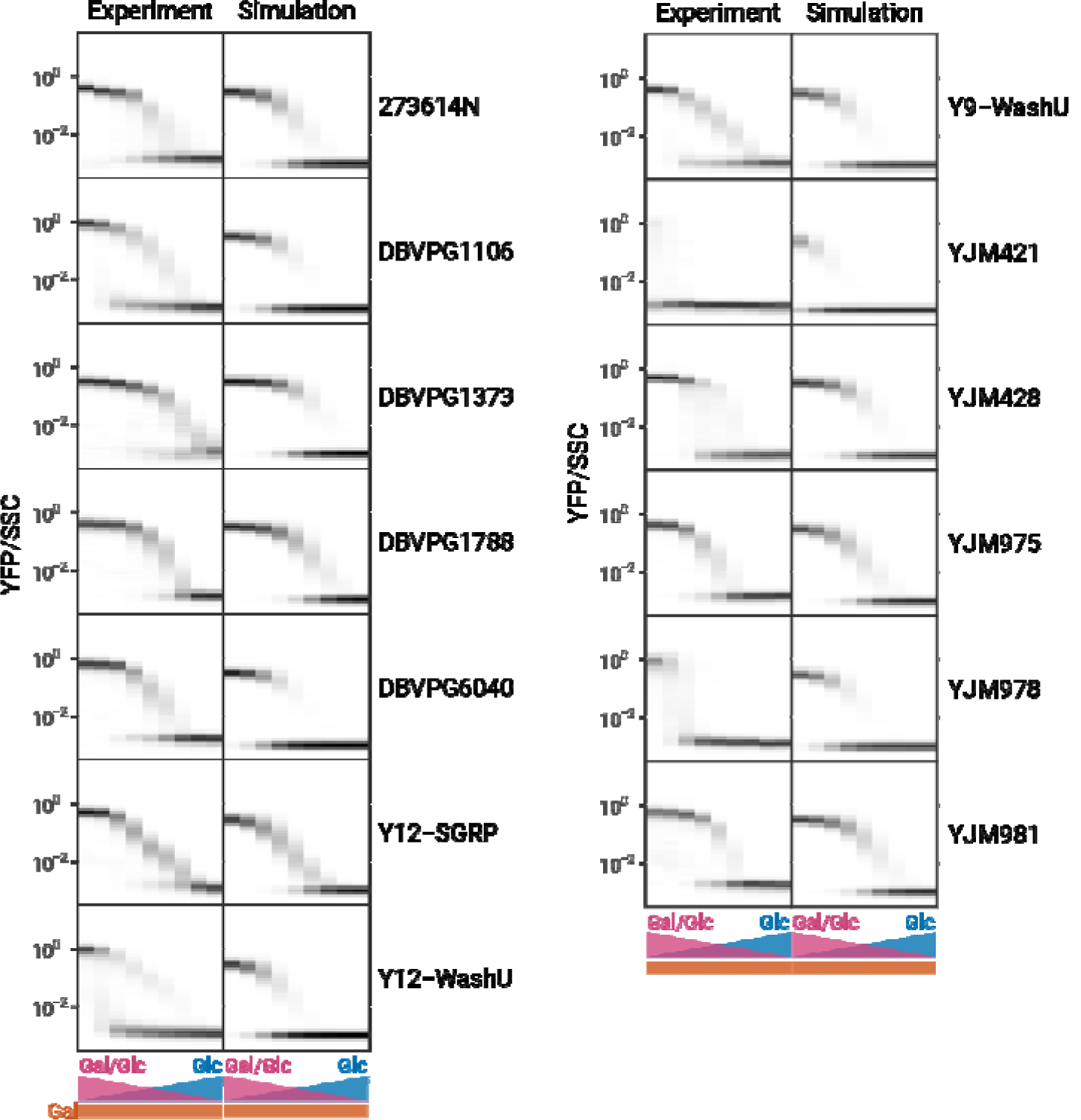
Experimental and simulated GAL induction profiles of bimodal strains. Simulations were performed as shown in Figure 3. Glucose concentration is titrated in two-fold steps from 1% to 0.0039%, galactose concentration is constant at 0.25%.

**Figure S7.**
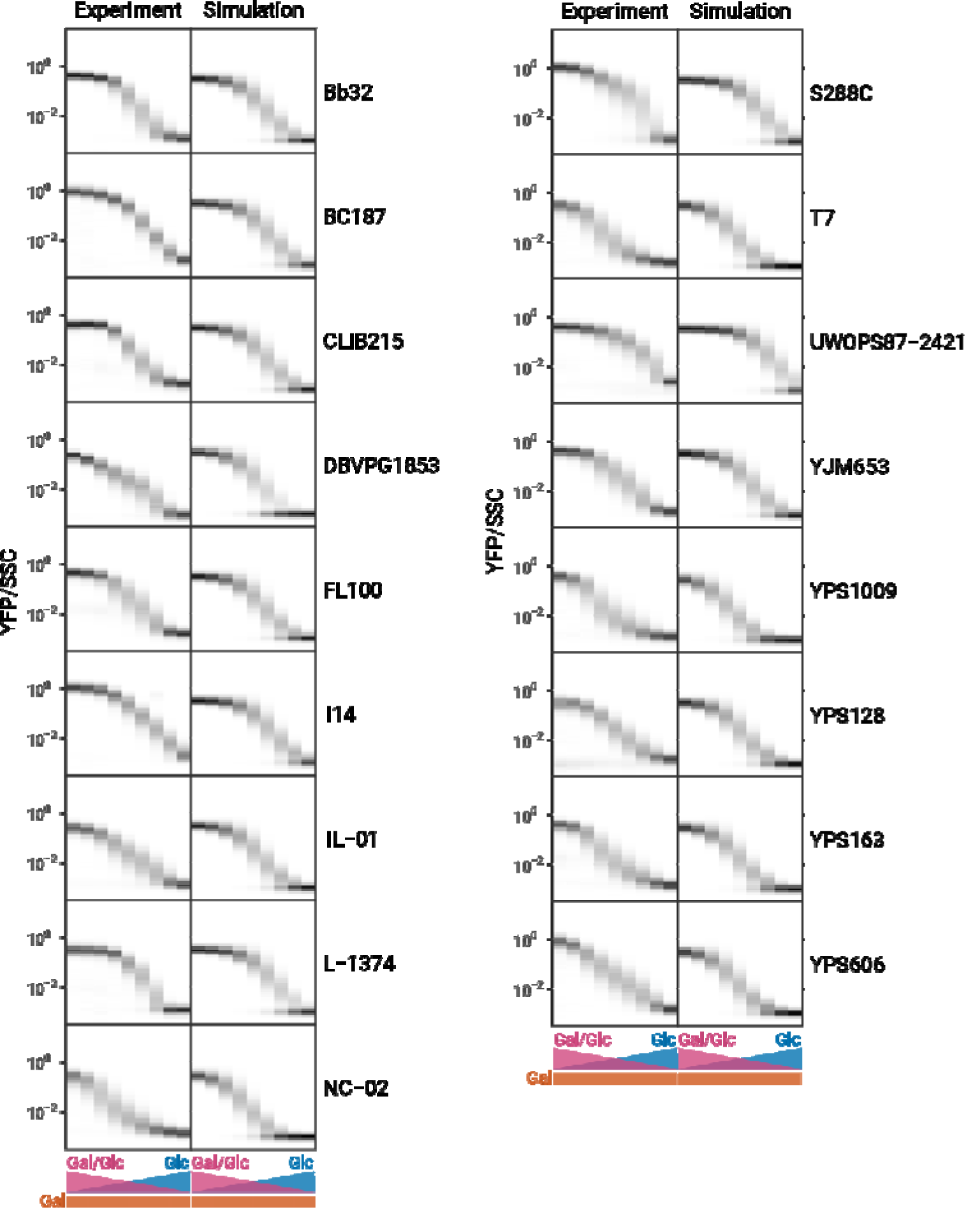
Experimental and simulated GAL induction profiles of unimodal strains. Simulations were performed as shown in Figure 3. Glucose concentration is titrated in two-fold steps from 1% to 0.0039%, galactose concentration is constant at 0.25%.

**Figure S8.**
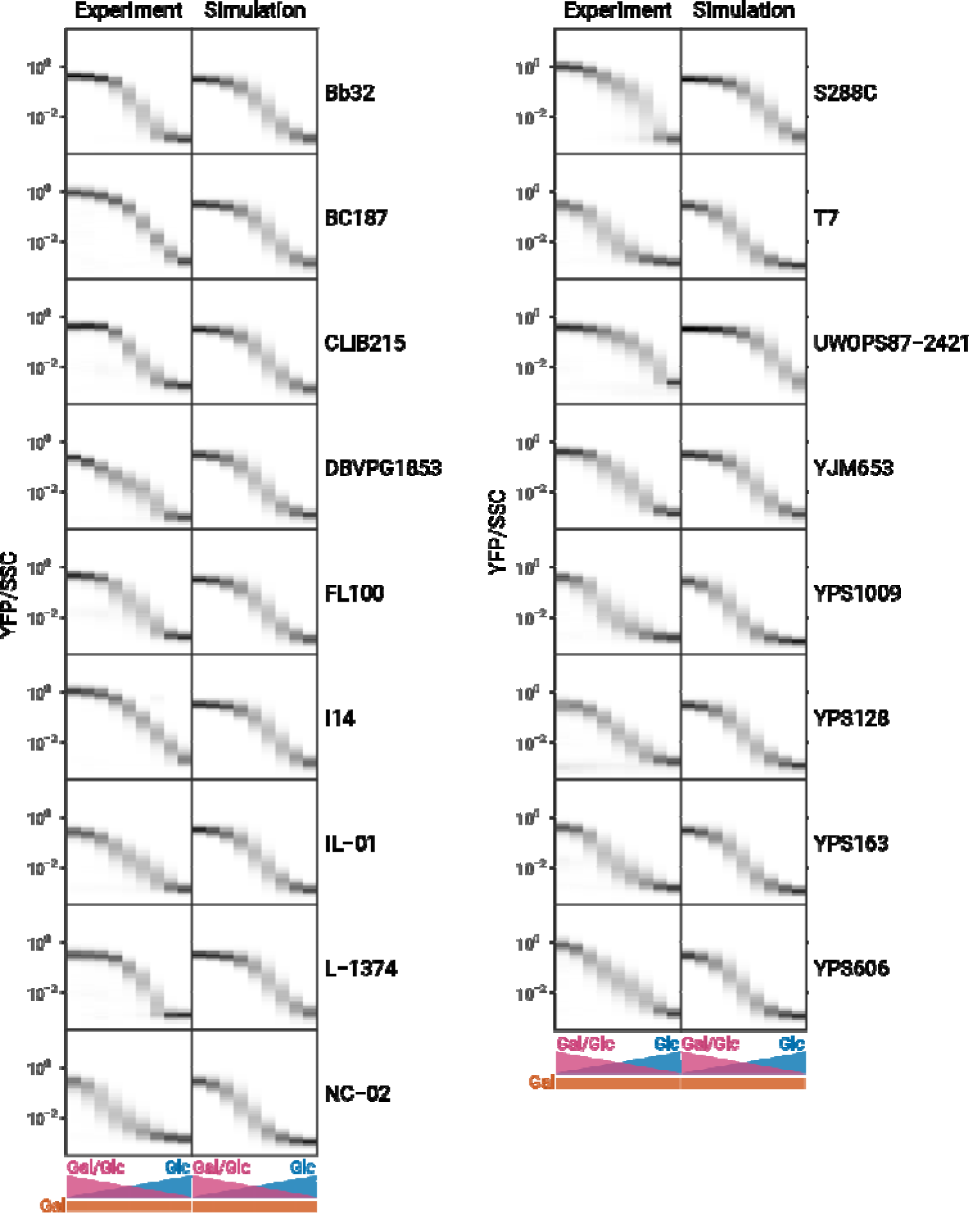
Experimental and simulated GAL induction profiles of unimodal strains with higher F_90_. Simulations were identical to those in Figure S6, but the experimentally determined F90 was increased two-fold for simulation purposes.

**Figure S9.**
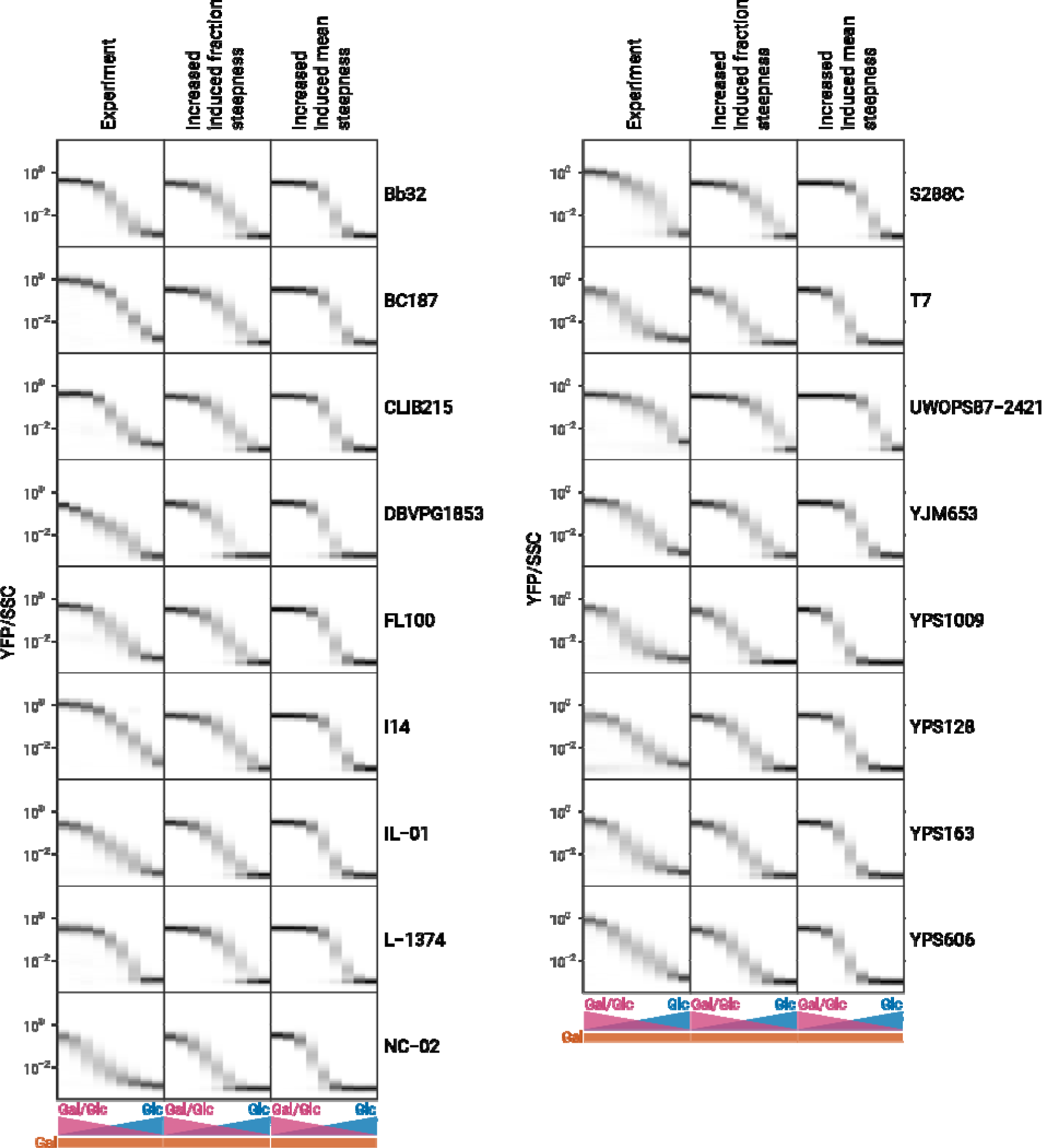
Experimental and simulated GAL induction profiles of unimodal strains with varying steepness. Simulations were identical to those in Figure S6, but the n parameter of the functions (see Figure 3 and Methods) was increased to the highest value that was observed when the functions were fitted to experimental data (induced level curve: 1.72, induced fraction curve: 3.10, see Figure S10).

**Figure S10.**
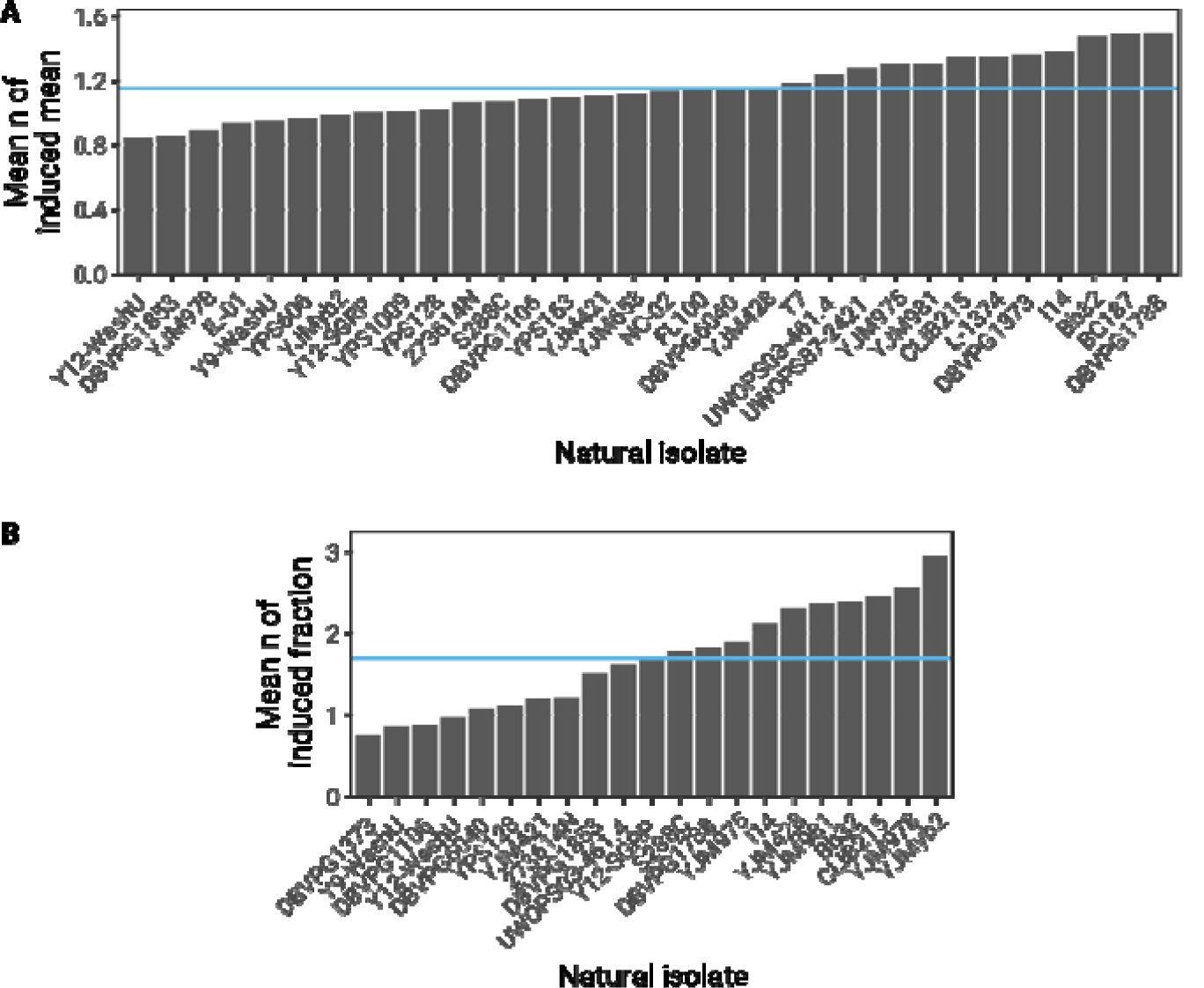
Fitted values for the n parameter. The functions described in Figure 3 were fitted to (A) the induced level curves of all strains or (B) the induced fraction curves of bimodal strains and the n parameter of the fit was extracted. Shown values represent the mean of fits to replicates. Blue lines indicate the mean of all means.

**Figure S11.**
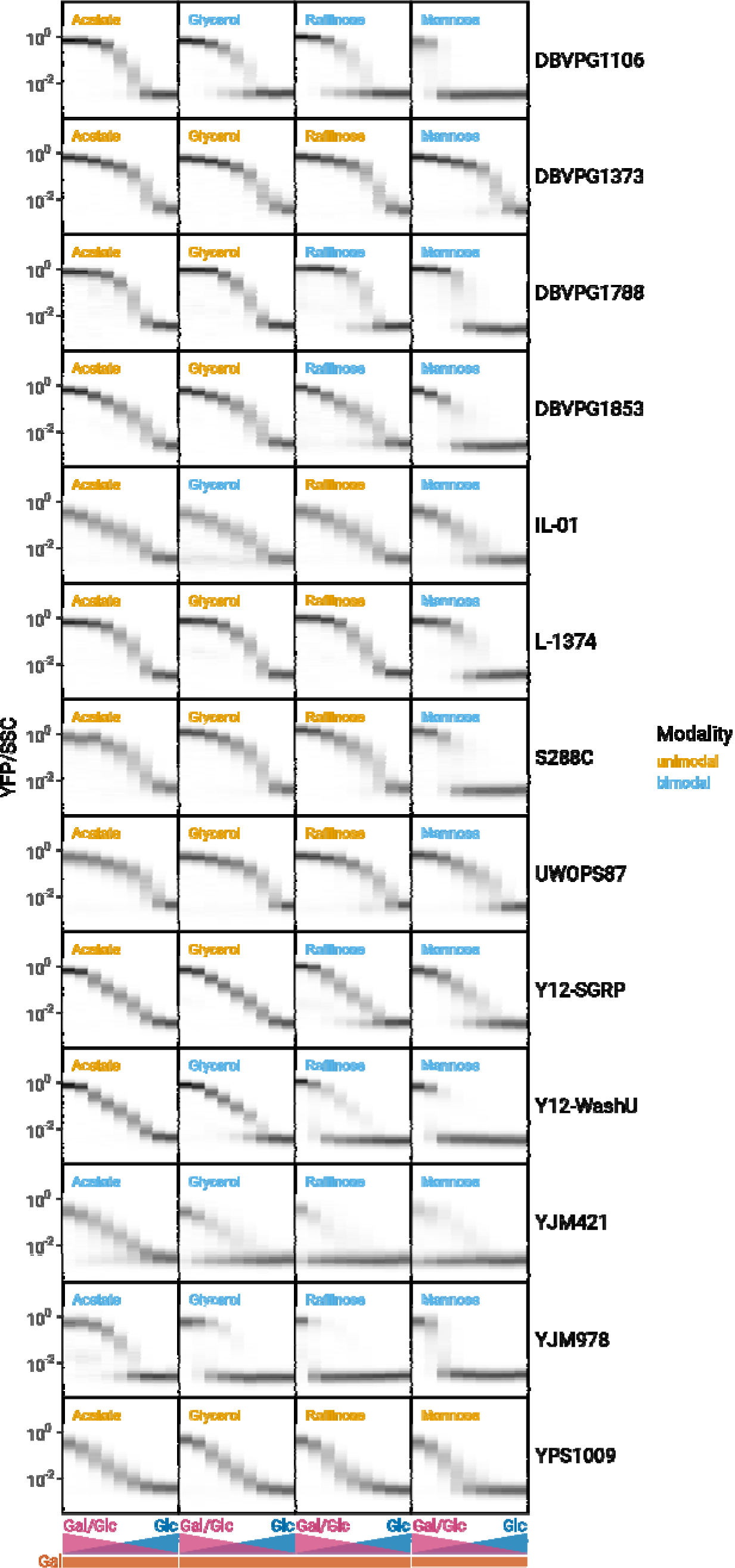
GAL induction of natural isolates in different glucose and galactose concentrations after growth in different carbon sources. Each plot represents 9 histograms with color intensities corresponding to the density of cells with a given fluorescence value (normalized by side-scatter (SSC)). Glucose concentration is titrated in two-fold steps from 1% to 0.0039%, galactose concentration is constant at 0.25%. Orange titles indicate unimodal induction, blue titles indicate bimodal induction.

**Figure S12.**
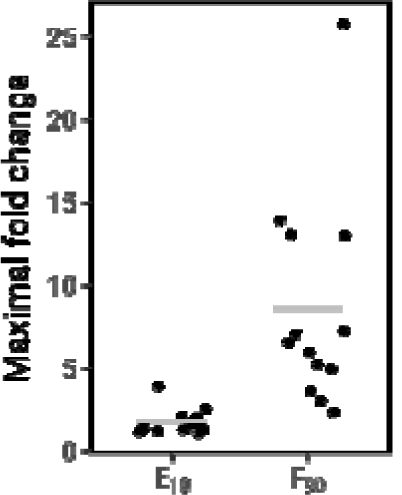
Fold change between the highest and lowest E10 and F90 values after growth in different pre-induction conditions for all isolates shown in Figure S11. Gray bars indicate the means of the maximal fold changes.

**Figure S13.**
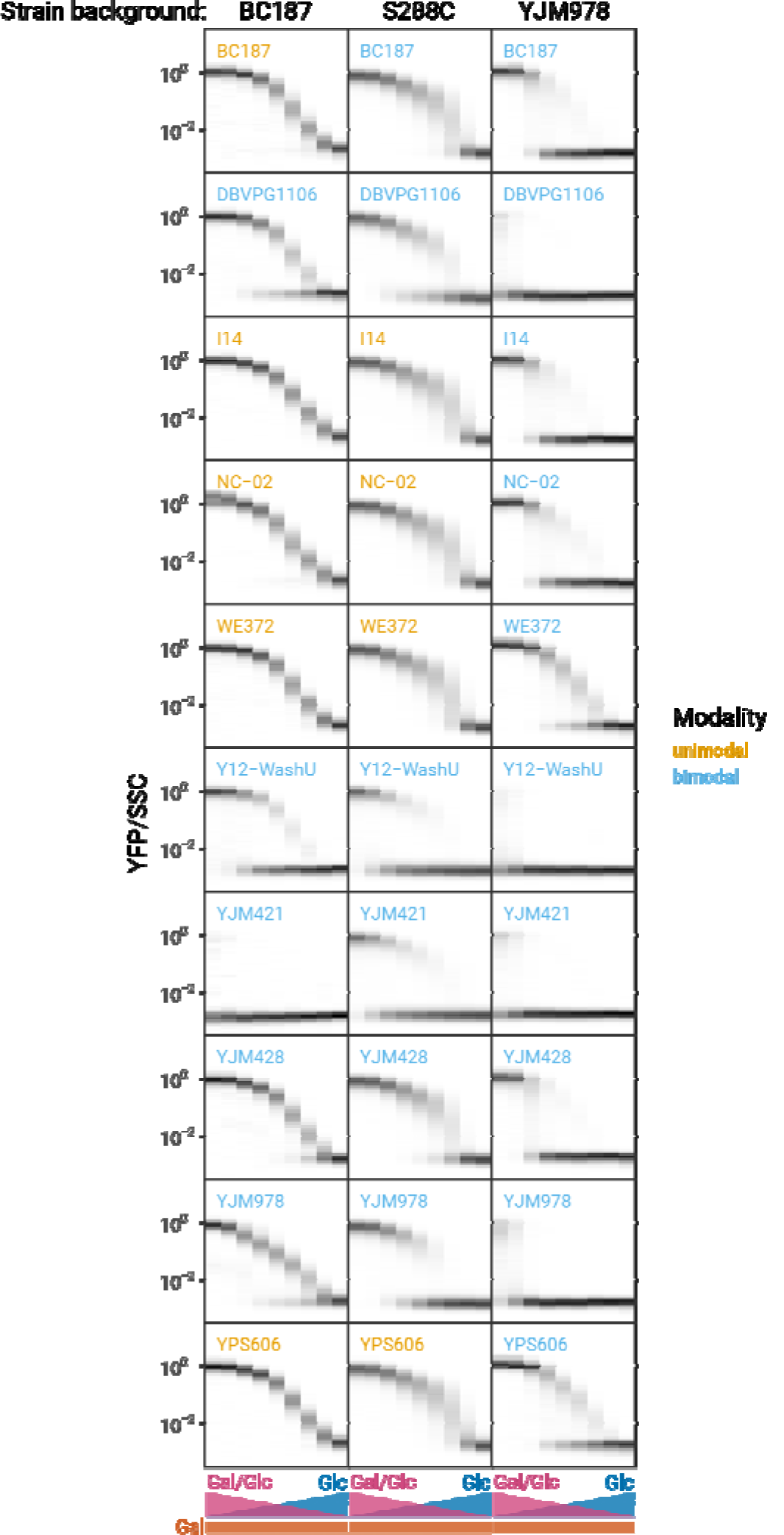
GAL induction of allele swap strains in different glucose and galactose concentrations. Each plot represents 9 histograms with color intensities corresponding to the density of cells with a given fluorescence value (normalized by side-scatter (SSC)). Glucose concentration is titrated in two-fold steps from 1% to 0.0039%, galactose concentration is constant at 0.25%. Plot titles indicate the source of the GAL3 allele. Orange titles indicate unimodal induction, blue titles indicate bimodal induction.

